# Introducing a translationally relevant mouse model of radiosurgery-induced unilateral hearing loss

**DOI:** 10.64898/2025.12.23.696156

**Authors:** Dimitrios Daskalou, Francis Rousset, Stéphanie Sgroi, Lucie Oberhauser, Jean-Philippe Thiran, Constantin Tuleasca, Ileana O Jelescu, Marc Levivier, Pascal Senn

**Affiliations:** The inner ear and olfaction lab, University of Geneva, Faculty of Medicine, Geneva, Switzerland; Service of Otorhinolaryngology-Head and Neck Surgery, Department of Clinical Neurosciences, Geneva University Hospitals, Geneva, Switzerland; Laboratory of Signal Processing 5, École Polytechnique Fédérale de Lausanne (EPFL), Lausanne, Switzerland; Department of Radiology, Lausanne University Hospital (CHUV), Lausanne, Switzerland; Neurosurgery Service and Gamma Knife Center, Centre Hospitalier Universitaire Vaudois (CHUV), Rue du Bugnon 44-46, BH-08, CH-1011 Lausanne, Switzerland; Hôpital de La Tour, Centre NeuroKnife, Av. J.-D.-Maillard 3, 1217 Meyrin (Geneva), Switzerland

**Keywords:** Radiosurgery, Vestibular Schwannoma, Hearing loss, Cochlea, Mouse model, Auditory brainstem response, Hair cells

## Abstract

**Background:** Stereotactic radiosurgery (SRS) is widely used to treat vestibular schwannomas but may cause irreversible hearing loss due to cochlear toxicity. The underlying mechanisms are not fully understood, and no effective otoprotective therapies exist. We aimed to establish a mouse model that replicates the clinical pattern of radiation-induced hearing loss.

**Methods:** C57BL/6J mice (n=31) received 8 Gy (n=3), 16 Gy (n=5), 24 Gy (n=8), or 32 Gy (n=15) using the Leksell Gamma Knife ICON system. Targeting was based on the built-in cone-beam CT, co-registered with MRI and CT-based mouse atlases, to guide unilateral cochlea targeting. A single isocenter was placed lateral to the right cochlea, with the 80% isodose line traversing its medial edge. Auditory brainstem response (ABR) was measured at baseline and at 1 and 4 weeks post-SRS. A 32 Gy subgroup (n=7) was evaluated at 16 weeks. Histological analysis of cochleae was performed at 4 weeks in all groups and at 16 weeks in the long-term 32 Gy group.

**Results:** SRS was well tolerated, and the contralateral cochlea received a very low radiation dose. No ABR shifts were observed at 8 or 16 Gy, with only minimal histological changes. At 32 Gy, ABR threshold shifts at 22.6 and 32 kHz were evident by week 1 and worsened by week 4. Similar but milder effects occurred at 24 Gy. In the 32 Gy long-term subgroup, hearing loss progressed across all frequencies, most severely at high frequencies, alongside a sustained wave I amplitude decline. At 32 Gy, outer hair cells were reduced by 14% and 44% at 32 and 45.2 kHz, respectively, at 4 weeks, and by 38% and 80% at 16 weeks. Ribbon synapses were mildly reduced at 4 weeks and more markedly at 16 weeks in corresponding high-frequency regions. Spiral ganglion neuron density was mildly reduced at the basal and middle turns. All reported changes were statistically significant when compared to the contralateral ear.

**Conclusions:** This new model reproduces key features of SRS-induced cochlear toxicity, including unilateral, dose-dependent, and progressive hearing loss. It thus provides a valuable platform for investigating mechanisms and testing otoprotective strategies.

## 1. Introduction

A vestibular schwannoma (VS) is a benign tumor arising from Schwann cells of the vestibulocochlear nerve and accounts for 8% of intracranial neoplasms (1). Despite being benign, VS can lead to hearing loss, vertigo, tinnitus, and, in some cases, symptomatic compression of the facial nerve, trigeminal nerve, and brainstem (2). Upon radiological progression or the onset of symptoms, VS needs to be treated either with neurosurgical excision or by radiation therapy. Stereotactic radiosurgery (SRS) is a precise radiation technique that plays a cornerstone role in the treatment of small-to-medium VS, achieving 5-year tumor control rates of 90-99% (3). SRS is delivered using various devices, such as the Leksell Gamma Knife (LGK; Elekta AB, Stockholm, Sweden), which uses multiple cobalt-60 sources, and dedicated linear accelerator-based systems like CyberKnife (Accuray, Sunnyvale, CA, USA). These devices enable focused radiation delivery while minimizing exposure to surrounding healthy tissues (4).

Despite this precision and recent advancements in SRS protocols for VS, hearing loss remains a significant concern. Indeed, recent meta-analyses demonstrate that hearing loss after SRS for VS is progressive, with more than 40% of patients losing their serviceable hearing within the first 4-6 years and 80% at ten years or more (5, 6). Currently, there is no preventive or therapeutic agent for radiation-induced hearing loss, highlighting the need for the development and experimental testing of novel treatments.

Previous studies have investigated the radiobiological effects of radiation-induced hearing loss in animal models and explored potential radioprotective agents (7–18). In these studies, the authors used biological irradiators or clinical linear accelerators, which delivered broad-field or whole-head irradiation with limited spatial precision. As a result, these approaches yielded approximate radiation doses delivered to both cochleae, while the central auditory system pathways were equally irradiated. These models do not accurately reflect radiation delivery in clinical SRS for VS, where the steep radiation decay can affect the cochlea but not the central pathways. Furthermore, these broad-field radiation models induce significant side effects in treated animals, with reported aversion to food and subsequent weight loss due to radiation-induced damage to the pharyngeal mucosa (9).

Here, we aimed to establish a new mouse model that replicates the pattern of radiation-induced hearing loss observed in patients treated with SRS for VS. This model offers several advantages over previously described approaches. First, it induces dose-dependent functional and histological cochlear damage, enabling quantitative assessment of radiotoxic effects. Second, the use of stereotactic targeting spares the central auditory pathways, thereby more accurately replicating the clinical scenario of SRS for VS. Third, the contralateral cochlea remains unaffected and can serve as an internal control, reducing inter-animal variability and the number of animals required. Finally, the localized irradiation protocol avoids damage to adjacent structures, such as the pharyngeal mucosa, thus preventing systemic side effects, in line with the 3Rs (replace, reduce, refine) principles.

## 2. Materials and methods

### 2.1. Animal model and ethics

All procedures involving animals were conducted in accordance with Swiss federal and cantonal animal welfare legislation and were approved by the Federal Food Safety and Veterinary Office (authorization number 36005) and the Cantonal Veterinary Office of Geneva (authorization number GE316).

C57BL/6J mice (n = 31), six weeks of age at baseline, were housed in groups under standard laboratory conditions (12-hour light/dark cycle, constant temperature and humidity) with *ad libitum* access to food and water. Animals were allowed a minimum of one week of acclimatization prior to baseline testing.

Experimental procedures involving potential discomfort, including auditory testing and SRS, were performed under general anesthesia induced by an intraperitoneal injection of ketamine (100 mg/kg; Ketasol, Graeub) and xylazine (10 mg/kg; Rompun, Elanco), prepared in sterile saline at a total volume of 10 μL/g body weight. Anesthesia depth was monitored regularly via pedal withdrawal reflex, and when necessary, a supplemental intramuscular injection of ketamine (100 mg/kg, 5 μL/g body weight) was administered to extend anesthesia duration. An ophthalmic ointment was applied during general anesthesia to ensure corneal protection.

Animal well-being was assessed daily during the first post-SRS week and then weekly, with body weight monitored regularly as an additional welfare parameter. No signs of distress, weight loss, or behavioral abnormalities were observed in any of the experimental groups.

The study included two experimental arms: a dose–response group (n = 24), in which animals received a dose of 8, 16, 24, or 32 Gy and were evaluated up to 4 weeks post-SRS; and a long-term follow-up group (n = 7), in which animals receiving 32 Gy were monitored up to 16 weeks post-SRS to assess the progression of auditory and histological damage.

To determine the optimal dose for inducing measurable hearing loss at week 4, we employed a stepwise, adaptive design. Based on the range of doses typically received by the cochlea in clinical VS SRS, we initially irradiated three mice with 8 Gy. As no auditory brainstem response (ABR) threshold shifts were observed, we followed the approved protocol by discontinuing this group and introducing a 16 Gy group. After treating five mice, only minimal functional and histological changes were observed, and detecting statistically significant effects at this dose would have required a prohibitively large number of animals. We therefore ceased further experiments at 16 Gy and focused on higher-dose groups (24 Gy and 32 Gy), which consistently produced measurable cochlear toxicity. This adaptive approach ensured compliance with ethical limits on animal use, allowed real-time refinement of the protocol, and supported the development of a robust and translationally relevant model.

At the study endpoint (week 4 and week 16 post-SRS), mice were euthanized under deep anesthesia by decapitation, in accordance with approved protocols. This method ensured rapid death without prior tissue fixation, enabling downstream cochlear histology and molecular analysis.

### 2.2. Stereotactic radiosurgery setup, imaging, and targeting

SRS was performed using the LGK ICON system (Elekta AB, Stockholm, Sweden), a clinical-grade SRS platform. Mice were anesthetized with ketamine and xylazine as previously described and placed in a prone position on a custom-designed stereotactic frame made from thermoplastic material (Fig. 1A). This frame was specifically designed to fit within the LGK system and was positioned at the patient’s head location on the robotic treatment bed. The frame was stabilized using a Moldcare cushion, ensuring secure immobilization and precise repositioning (Fig. 1B).

**Fig 1.**
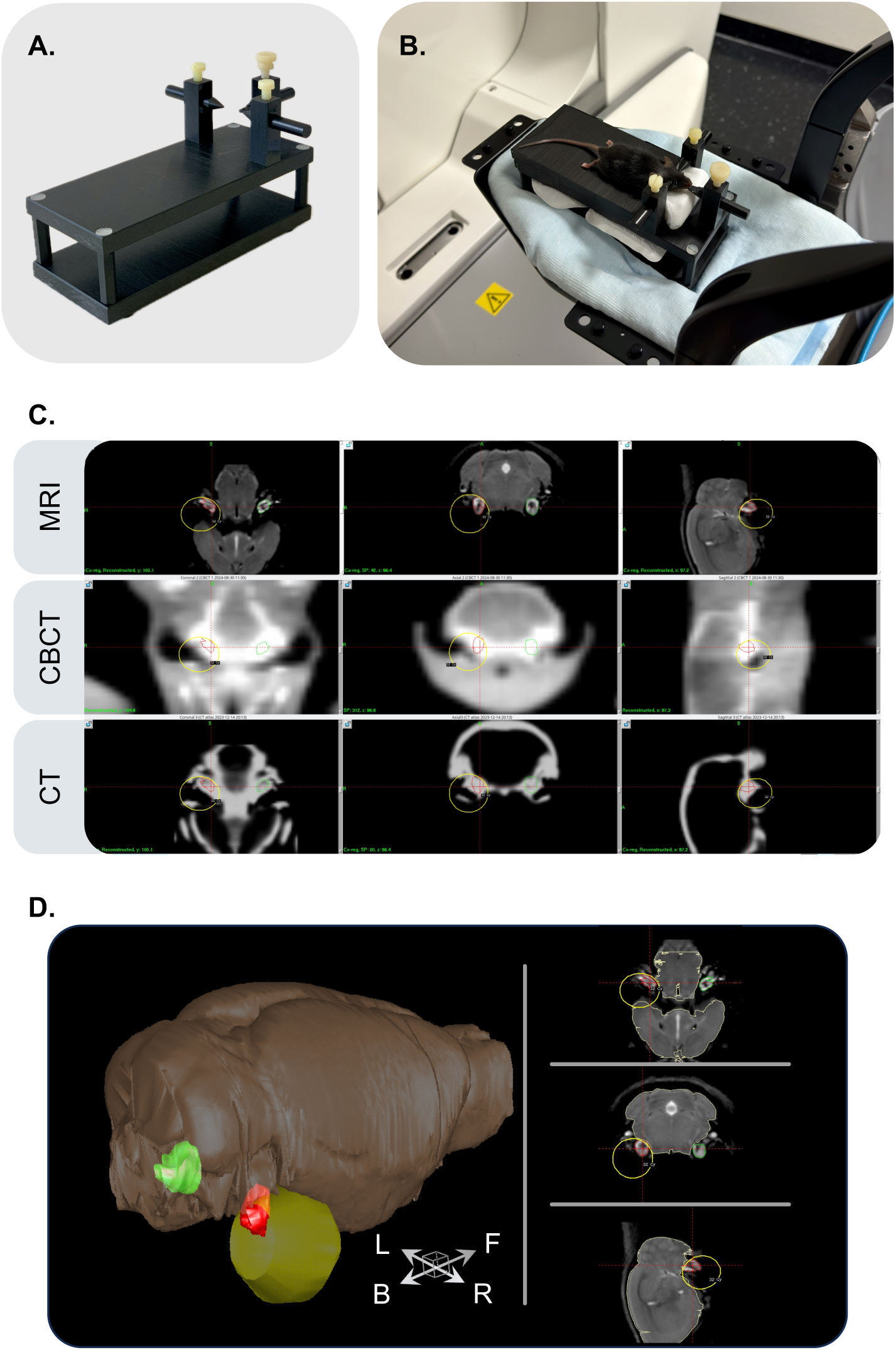
Stereotactic radiosurgery setup and image-guided targeting protocol. (A) Custom-designed stereotactic frame used to position and immobilize mice in prone orientation on the Gamma Knife treatment couch. (B) Immobilization system stabilized with a Moldcare cushion to ensure accurate repositioning at the head support location of the Gamma Knife treatment couch. (C) Cone-beam computed tomography (CBCT)-based irradiation planning. The upper row corresponds to 3 planar imaging from a mouse atlas magnetic resonance imaging (MRI), and the lower row to high-resolution computed tomography (CT) mouse atlas. The middle row corresponds to a representative example of a Gamma Knife-acquired CBCT of an experimental animal used to plan targeted irradiation. The three image modalities are co-registered and processed in Leksell Gamma Plan, the dosimetry software of the Gamma Knife system. The right cochlea is contoured in red and the contralateral cochlea in green. A single isocenter is placed lateral to the right cochlea. The 80% isodose line (yellow line) passes through the medial cochlear edge, and in this example, corresponds to a dose of 32 Gy. (D) MRI-based 3D reconstruction after co-registration of CBCT with MRI and CT mouse atlas to delineate cochlear structures and perform volumetric segmentation for dose-volume analysis. The right irradiated cochlea is depicted in red, the left contralateral cochlea is shown in green, and the yellow sphere represents the 80% isodose line volume, corresponding to a dose of 32 Gy. The brown-grey volume corresponds to the brain, based on the MRI atlas segmentation.

An initial built-in cone-beam computed tomography (CBCT) scan was acquired to visualize each animal’s skull and cochlear anatomy. To ensure precise anatomical targeting of the cochlea, CBCT images acquired with the LGK ICON system were co-registered with a CT-based anatomical atlas of the mouse head, followed by a high-resolution magnetic resonance imaging (MRI) mouse brain atlas (19, 20) (Fig. 1C). Bilateral segmentation of the cochleae was performed using the co-registered images to create individualized three-dimensional volumes of interest (Fig. 1D). Cochlear segmentation and dose–volume analysis were performed using the volumetric tools within Leksell GammaPlan (LGP; Elekta AB, Stockholm, Sweden). The resolution of the CBCT scans was sufficient to resolve the cochlear contours, and the co-registered MRI provided additional anatomical confidence, particularly in defining medial and lateral cochlear borders for isocenter placement.

Radiation treatment planning was performed in LGP software, in which the specific CBCT was co-registered with imported MRI and CT-based mouse atlases, allowing accurate placement of a single 4 mm isocenter lateral to the right cochlea (Fig. 1C). The isocenter was positioned so that the 80% isodose line was aligned to pass through the medial edge of the cochlea, a configuration selected to replicate clinical conditions in SRS for VS, where the cochlea lies in the high-gradient periphery of the radiation field. This setup enabled unilateral irradiation while sparing the contralateral cochlea and minimizing off-target exposure to surrounding structures, including the brainstem and pharyngeal mucosa.

Each animal was assigned to receive a single-fraction dose of 8 (n = 3), 16 (n = 5), 24 (n = 8), or 32 (n = 15) Gy prescribed to the 80% isodose. Following plan approval, a second CBCT scan was performed to detect and correct any head displacement (as per the obligatory clinical workflow of the frameless LGK ICON procedure), although no corrections were necessary with animals under general anesthesia. Subsequently, the robotic couch automatically adjusted to align the target with the planned isocenter. The full procedure, including positioning, imaging, plan verification, and radiation delivery, was completed within approximately 20–30 minutes per animal, depending on the radiation dose.

To maintain normothermia during anesthesia, air-activated heating pads were placed beneath the thermoplastic base supporting the animal. These pads generate heat through a controlled exothermic oxidation reaction of iron powder, providing consistent thermal support without the need for electrical components that could interfere with the LGK system. Before starting the study, we characterized the temperature profile of the pads and established a replacement interval to ensure stable heating conditions throughout the procedure.

This image-guided workflow enabled accurate dose-volume calculations, reproducible near-cochlear targeting across animals, and minimized inter-individual variability in radiation delivery. The spatial targeting achieved by this protocol, including the relative positioning of the cochleae, brain structures, and the 80% isodose line, is illustrated in Figures 1C and 1D.

## 2. 3. Auditory brainstem response testing

Hearing thresholds were assessed by ABR recordings one day before SRS (baseline), at 1 and 4 weeks post-SRS in the dose–response groups, and at 4, 8, 12, and 16 weeks post-SRS in the long-term follow-up group. All measurements were performed in a sound-attenuated chamber (IAC Acoustics, IL, United States) with animals under general anesthesia, as described in Section 2.1. Throughout the procedure, animals were placed on an isothermal pad to maintain normothermia.

ABRs were recorded from both ears independently using custom-fitted ear adapters to direct acoustic stimuli into each ear canal, with the contralateral ear serving as an internal control. Subdermal electrodes were positioned at the forehead (active), the mastoid of the stimulated ear (reference), and the back of the experimental animal (ground). Acoustic stimuli were generated and responses recorded using a multifunction IO card (National Instruments, Austin, TX, USA). Sound output was calibrated online prior to each recording using a microphone probe system (Brüel & Kjær 4191) placed near the animal’s ear. Signal amplitude was adjusted using an attenuator and amplifier (Otoconsult, Frankfurt, Germany), and recordings were amplified (80 dB) and band-pass filtered (0.2–3.0 kHz).

Each session included ABRs evoked by both click stimuli (100 µs broadband pulse) and nine pure tone bursts (ranging from 2 to 32 kHz, in two steps per octave; 3 ms duration). Each stimulus was presented 128 times, and responses were averaged to enhance signal clarity. Click stimuli were delivered in 3 dB sound pressure level (SPL) increments and tone bursts in 5 dB SPL increments, ranging from 0 to 90 dB SPL. ABR thresholds were determined as the lowest sound intensity at which a consistent and visually detectable wave pattern was observed.

The wave I amplitude was measured as the voltage difference between the first positive peak and the subsequent negative trough in response to click stimuli. Values were averaged across suprathreshold intensities ranging from 72 to 78 dB SPL.

After each recording session, animals were returned to their home cages and monitored on a heating pad until full recovery from anesthesia.

## 2. 4. Cochlear dissection and preparation

Following the final ABR measurement at week 4 and week 16, mice were euthanized. Temporal bones were rapidly extracted to minimize post-mortem tissue degradation. A sagittal incision was made along the skull, and both temporal bones were harvested. Under a stereomicroscope, the bony auditory bulla was dissected to expose the cochlea, which was subsequently transferred to cold PBS (1X, Gibco). Upon opening the middle ear, we confirmed the absence of fluid in all of the experimental animals. Care was taken to preserve the structural integrity of the cochlea, and no specimens showed mechanical damage.

Cochleae were fixed in 4% paraformaldehyde (PFA) at 4 °C overnight. Decalcification was performed using a commercial decalcification reagent (USEDECALC, Art. No. 40-3310-00) at room temperature for 48 hours, with a solution change after 24 hours. After decalcification, both cochleae from each mouse were randomly assigned to one of two downstream applications: (1) paraffin embedding for mid-modiolar sectioning, or (2) whole-mount dissection followed by immunofluorescent analysis of the sensory epithelium.

## 2. 5. Cochlear histology

## 2. 5. 1. Whole-mount immunohistochemistry and confocal microscopy

Decalcified cochleae assigned for cytocochleograms were dissected following the technique previously described (21). The epithelium was divided into three segments corresponding to the basal, middle, and apical cochlear turns. These segments were adhered to 12 mm round coverslips using Cell-Tak tissue adhesive (Corning, Cat# CLS354240), as described by Fang et al. (22), and then placed in PBS in a 24-well plate and stored at 4°C until immunostaining (Supplementary Fig. S1A-C).

Tissue samples were permeabilized with 2% Triton X-100 in PBS for 30 minutes at room temperature, followed by incubation in a blocking buffer containing 2% bovine serum albumin and 0.01% Triton X-100 in PBS for 30 minutes. Primary antibody incubation was performed overnight at 4 °C using rabbit anti-Myosin VIIa (1:200; Proteus Biosciences, Cat# 25-6790) and mouse anti-CtBP2 (1:200; BD Biosciences, Cat# 612044), both diluted in blocking buffer. After three PBS rinses, samples were incubated for 2 hours at room temperature with donkey anti-rabbit Alexa Fluor 488 (1:500; Thermo Fisher Scientific, Cat# A21206) and donkey anti-mouse Alexa Fluor 555 (1:500; Thermo Fisher Scientific, Cat# A31570). Tissues were washed and mounted on glass slides with Fluoroshield mounting medium containing DAPI (Sigma, Cat# F6057).

Image acquisition was performed on a Zeiss LSM800 confocal microscope. For hair cell quantification, Z-stacks were acquired using a Plan-Apochromat 20x/0.8 objective (pixel size: 0.624 µm, step size: 3 µm). The Tiles function was used to image adjacent fields, and images were automatically stitched to reconstruct the full cochlear segment. Representative images for figures were taken using a Plan-Apochromat 20x/0.8 objective (pixel size: 0.156 µm, step size: 0.49 µm). For ribbon synapse analysis, Z-stacks spanning 10–15 µm were acquired with a Plan-Apochromat 63x/1.4 oil immersion objective (pixel size: 0.099 µm, step size: 0.5 µm). All images were processed and analyzed using Fiji (ImageJ).

## 2. 5. 2. Mid-modiolar hematoxylin and eosin staining

Cochleae allocated for paraffin embedding were dehydrated in a graded ethanol series and embedded in paraffin. Serial mid-modiolar sections of 5 µm thickness were prepared and mounted on glass slides. Hematoxylin and eosin (H&E) staining was performed using Harris’ hematoxylin (5 min) followed by differentiation in acid alcohol, and a brief counterstain with eosin (2–3 s). Slides were dehydrated, cleared in xylene, and coverslipped using Eukitt mounting medium.

Image acquisition was performed using a Zeiss Axio Scan.Z1 slide scanner using a Plan-Apochromat 20x/0.8 objective (pixel size: 0.22 µm). Image analysis of spiral ganglion neuron (SGN) density and stria vascularis thickness was performed using QuPath software.

## 2. 5. 3. Quantitative analysis of cochlear morphology

Hair cell survival was assessed from cytocochleograms acquired using the Tiles and Z-stack functions of a Zeiss LSM800 confocal microscope with a 20x objective. Multiple z-stack tile images were obtained for each cochlear turn and stitched using ZEN Blue software to reconstruct the full extent of each turn. Z-stacks were then maximum-intensity projected, and the apical, middle, and basal turns were combined into a single image. Using Fiji (ImageJ) and the Measure_Line plugin from the Eaton-Peabody Laboratories Histology Core at Mass Eye and Ear (https://masseyeandear.org/research/otolaryngology/eaton-peabody-laboratories/histology-core), a line was drawn along the lateral margin of the inner hair cell (IHC) row to measure the total length of the cochlear sensory epithelium. This line also generated a mask corresponding to the place-frequency map, which was used to define frequency-specific regions of interest. IHC and outer hair cells (OHC) were then manually counted within 300 µm-long segments corresponding to each mapped frequency (Supplementary Fig. S1D).

Ribbon synapse quantification was performed on confocal Z-stacks of the Organ of Corti (10–15 µm depth, 0.5 µm steps) projected into single-plane images. CtBP2-positive puncta were manually counted within a span of 10–15 IHCs at defined positions along the previously generated frequency map, allowing frequency-specific ribbon density analysis (Supplementary Fig. S1E). The total number of puncta was then divided by the number of IHC in the corresponding region to calculate the average number of puncta per IHC.

SGN density was assessed from four H&E-stained non-consecutive mid-modiolar sections for each cochlea. The Rosenthal’s canal was manually annotated in QuPath and categorized by cochlear region (apex, middle, base). Using the Cellpose extension and a custom analysis script, the canal area and SGN nuclei within each region were automatically quantified. Automated results were subsequently validated by manual review to ensure accuracy. Counts were normalized to the annotated area (SGN/mm²), averaged per cochlear turn within each ear, and analyzed for group comparisons.

Stria vascularis thickness was assessed on H&E-stained mid-modiolar cochlear sections using QuPath. In each cochlea, measurements were performed on four non-consecutive sections. For each section, a line was drawn from the outer margin of the stria vascularis to the junction between basal cells and the spiral ligament, positioned halfway between the attachment points of Reissner’s membrane and the spiral prominence (23). This approach was applied to the basal, middle, and apical turns of both the irradiated and contralateral cochleae. Mean stria vascularis thickness values were calculated per turn for statistical comparison.

## 2. 6. Statistical analysis

Two-way repeated measures ANOVA was used to compare the irradiated and contralateral cochleae, with within-subject factors such as treatment side (irradiated vs. contralateral), stimulus frequency, cochlear region, or time, depending on the outcome. Data were paired within animals. Bonferroni correction was applied for *post hoc* pairwise comparisons. GraphPad Prism version 10.4.2 (534) was employed for data analysis and figure creation. The statistical significance was defined as p < 0.05 (two-sided).

## 3. Results

### 3.1. Stereotactic radiosurgery achieves precise, unilateral cochlear targeting

Across all experimental groups, SRS dosimetry was successfully focused on the right cochlea while minimizing off-target exposure. The contralateral cochlea consistently received less than 15% of the mean dose delivered to the irradiated cochlea. Mean radiation doses to the non-targeted cochlea were 1.2 (standard deviation [SD] = 0.3) Gy, 2.42 (SD = 0.13) Gy, 3.6 (SD = 0.33) Gy, and 4.83 (SD = 0.44) Gy for the 8, 16, 24, and 32 Gy groups, respectively.

The irradiation procedure was well tolerated, with no signs of behavioral changes or systemic adverse effects unrelated to hearing observed during the 16-week follow-up period. All animals gained body weight as expected for the strain, and no mortality occurred. In two animals from the long-term (16-week) 32 Gy group, we noted a slight discoloration of the fur in the retro-mandibular region following a linear pattern, which was not associated with any observable discomfort or other pathology.

### 3.2. ABR threshold shifts at 4 weeks are dose-dependent and frequency-specific

In the 8 Gy and 16 Gy groups, no significant ABR threshold shifts were observed at any time point (Fig. 2A,B,E,F). In contrast, mice in the 24 Gy and 32 Gy groups exhibited dose-dependent and progressive hearing loss in the irradiated ear, particularly at high frequencies (Fig. 2I). More precisely, at week 1, the 32 Gy group showed early threshold shifts at 22.6 and 32 kHz, with a mean difference between irradiated and contralateral ears of 11.25 dB at both frequencies (p = 0.006 for both; Fig. 2D). In the 24 Gy group, no significant shifts were detected at this time point (Fig. 2C). By week 4, threshold shifts became more pronounced. The 24 Gy group showed mean interaural differences of around 12 dB from 8 to 32 kHz, while in the 32 Gy group, the threshold shift differences reached 25 dB at 22.6 kHz (p < 0.0001) and 25.6 dB at 32 kHz (p < 0.0001) (Fig. 2G,H).

**Fig. 2.**
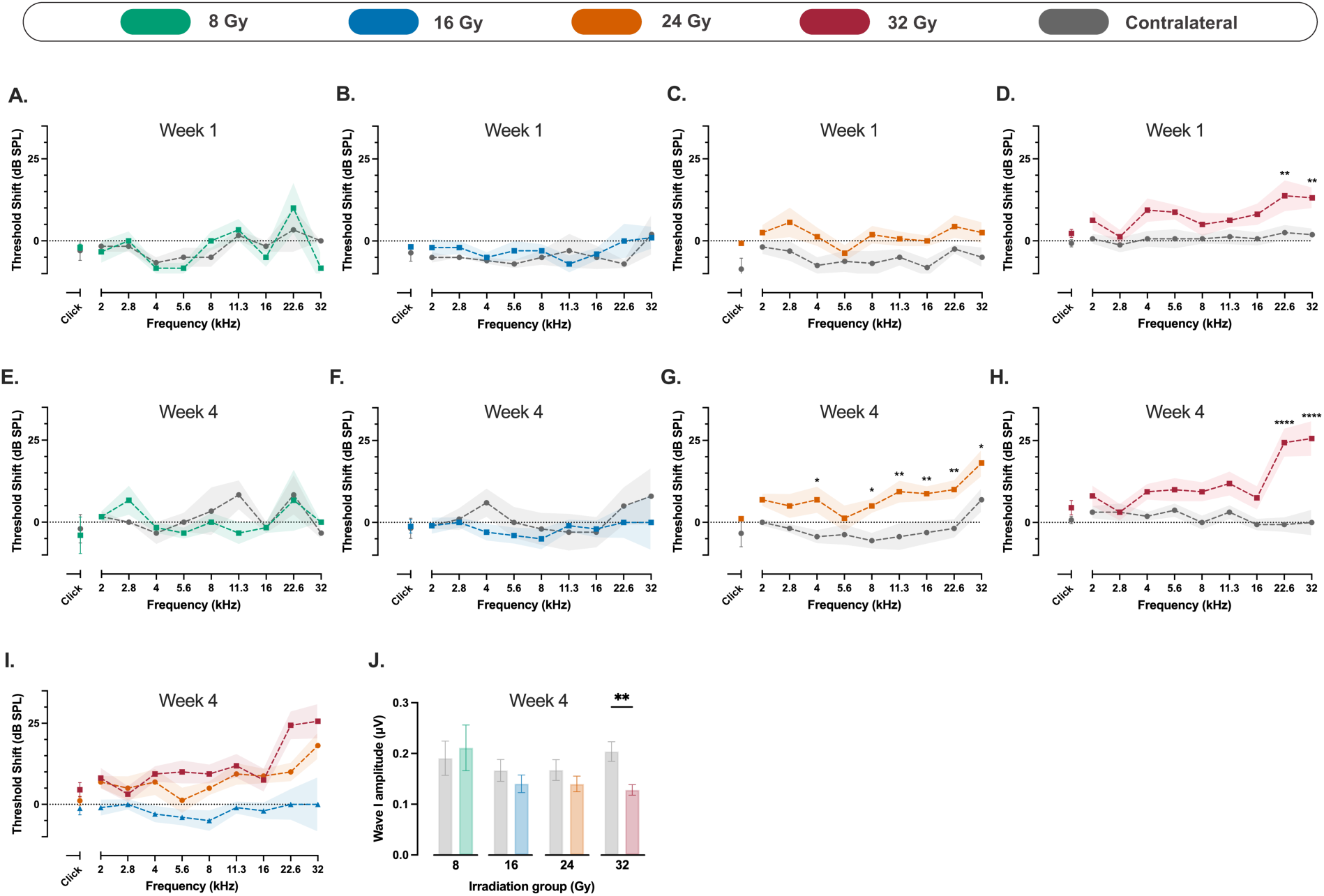
Auditory brainstem response (ABR) threshold shifts and wave I amplitude changes following unilateral cochlear irradiation. (A–H) Mean ABR threshold shifts (difference between baseline and indicated time point) in mice exposed to 8 Gy (A,E; n = 3), 16 Gy (B,F; n = 5), 24 Gy (C,G; n = 8), and 32 Gy (D,H; n = 8) at week 1 (A–D) and week 4 (E–H) post-irradiation. No significant threshold shifts were observed in the 8 Gy and 16 Gy groups at either time point. In contrast, mice receiving 24 Gy and 32 Gy showed progressive, dose-dependent hearing loss, with the most pronounced threshold shifts observed in the 32 Gy group at high frequencies (22.6 and 32 kHz) by week 4. (I) Mean ABR threshold shifts from the irradiated ear across 16, 24, and 32 Gy groups at week 4, highlighting dose-dependent effects. (J) Wave I amplitude of suprathreshold click-evoked ABRs at week 4 for the four experimental groups. A significant reduction in wave I amplitude was observed in the 32 Gy group in the irradiated ear compared to the contralateral ear. Data are presented as means with shaded areas or error bars representing SEM. Asterisks indicate statistical significance (*p < 0.05, **p < 0.01, ****p < 0.0001). Statistical analysis was performed using two-way repeated measures ANOVA with Bonferroni correction for *post hoc* pairwise comparisons. Colors correspond to radiation dose groups as shown in the legend. The contralateral ear serves as control in all cases.

We analyzed the amplitude of wave I after suprathreshold click stimulation. In the 16 and 24 Gy groups, a mild reduction in wave I amplitude was observed at week 4, but this difference compared to the contralateral ear did not reach statistical significance. In the 32 Gy group, at week 4, we observed a 37.1% lower amplitude at the irradiated ear compared to the contralateral side (mean difference, 0.08 μV; p = 0.003; 95% CI: 0.02 to 0.13; Fig. 2J). This reduction in wave I amplitude was already observed at week 1, albeit to a lesser extent. We did not find changes in the latency of wave I in all experimental groups (Supplementary Fig. S2A-D).

### 3.3. Progressive, irreversible hearing loss emerges in long-term follow-up

A subgroup of seven mice, irradiated with 32 Gy, was followed for an extended period of 16 weeks. ABR threshold shifts progressed over time and affected all measured frequencies at a significantly faster pace than expected for age-related hearing loss, as observed in the contralateral ear (Fig. 3A).

**Fig. 3.**
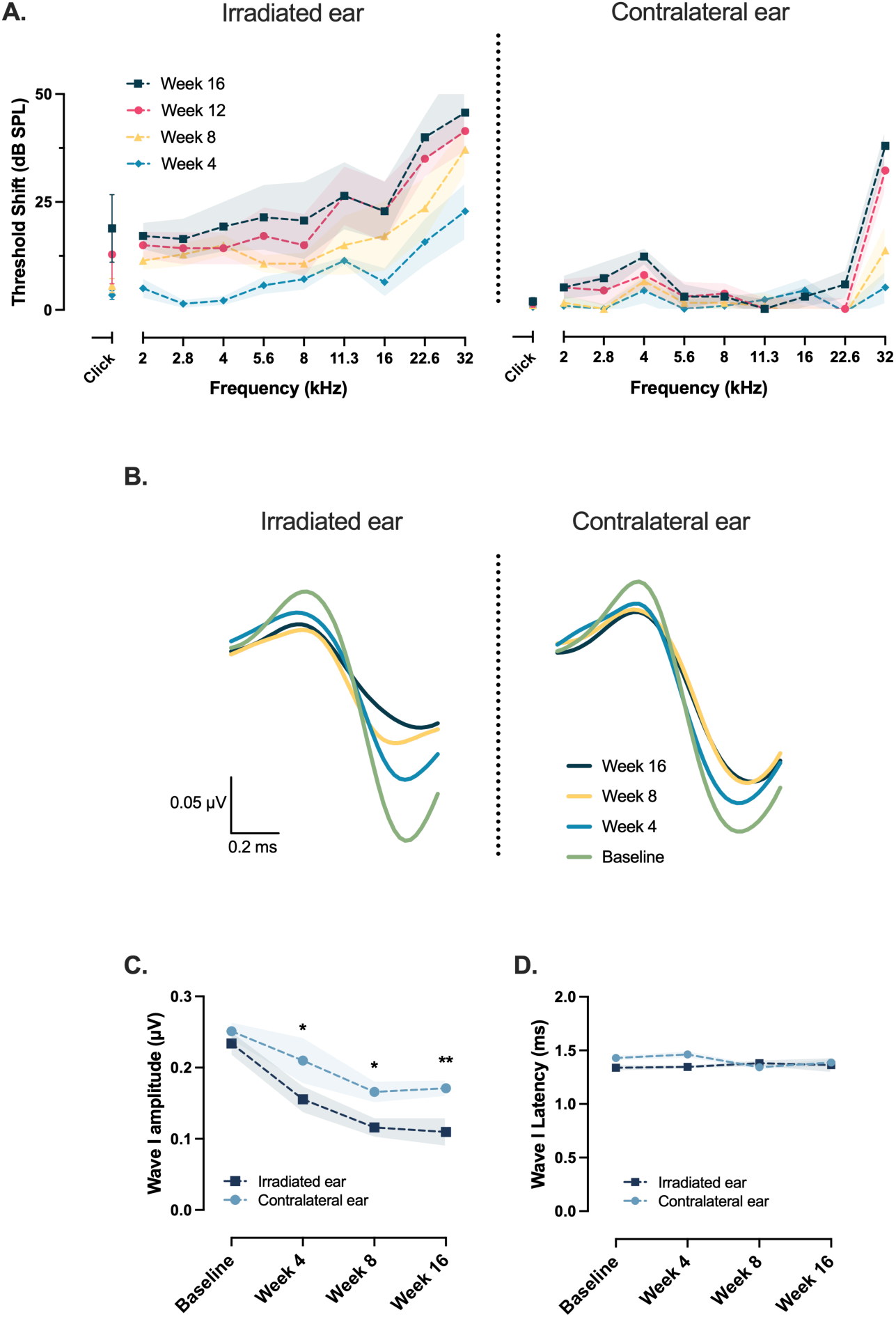
Auditory brainstem response (ABR) threshold shifts and wave I amplitude changes over a 16-week follow-up period after unilateral cochlear irradiation with 32 Gy (n = 7). (A) ABR threshold shifts (difference between baseline and each time point) progressively increased in the irradiated ear across all tested frequencies, whereas the contralateral ear showed only age-related shifts at high frequencies. (B) Averaged wave I waveforms. Wave I amplitude, in response to suprathreshold click stimulation, decreased progressively in the irradiated ear over time, while remaining relatively stable in the contralateral ear. (C) Quantification of wave I amplitude differences between irradiated and contralateral ears at baseline, 4, 8, and 16 weeks post-irradiation. (D) No significant changes were observed in wave I latency over time in either ear. Data are presented as means with shaded areas representing SEM. Asterisks indicate statistical significance (*p < 0.05, **p < 0.01). Statistical analysis was performed using two-way repeated measures ANOVA with Bonferroni correction for *post hoc* pairwise comparisons.

Consistent with threshold shift, we observed a progressive decline in wave I amplitude at the irradiated ear starting from week 4 and continuing to decrease further during the follow-up period (Fig. 3B). This decline was more pronounced compared to the natural age-related decline seen in the contralateral ear. More precisely, compared to the contralateral ear wave I amplitude on the irradiated side was on average 26% lower at 4 weeks (mean difference, 0.06 μV; p = 0.01; 95% CI: 0.01 to 0.1), 30% lower at 8 weeks (mean difference, 0.05 μV; p = 0.03; 95% CI: 0.005 to 0.1) and 36% lower at 16 weeks (mean difference, 0.06 μV; p = 0.005; 95% CI: 0.02 to 0.1; Fig. 3C). We did not find changes with time in the latency of wave I (Fig. 3D).

## 3. 4. Outer hair cell loss mirrors functional hearing decline

No significant loss of IHCs was observed in the irradiated cochleae across any of the four experimental groups (Supplementary Fig. S2E-H). In contrast, OHC loss showed a dose-dependent reduction in the high-frequency region corresponding to 45.2 kHz, assessed 4 weeks after SRS. Compared to the contralateral ear, OHCs at 45.2 kHz were reduced by 4.2 percentage points in the 8 Gy group (p = 0.01; 95% CI: 1.2 to 7.3; Supplementary Fig. S2I), 6.2 in the 16 Gy group (p = 0.003; 95% CI: 2.1 to 10.3; Supplementary Fig. S2J), 6.5 in the 24 Gy group (p = 0.002; 95% CI: 2.4 to 10.6; Fig. 4A), and 40.1 in the 32 Gy group (p < 0.0001; 95% CI: 25 to 55.1; Fig. 4B,C). A significant OHC loss was also observed at 32 kHz in the 32 Gy group, with a reduction of 13.3 percentage points compared to the contralateral cochlea (p = 0.03; 95% CI: 2.2 to 24.3). These findings confirm that OHCs in the basal cochlear turn are particularly susceptible to radiation, with greater damage at higher doses and frequency regions, paralleling the functional deficits observed in ABR threshold shifts.

**Fig. 4.**
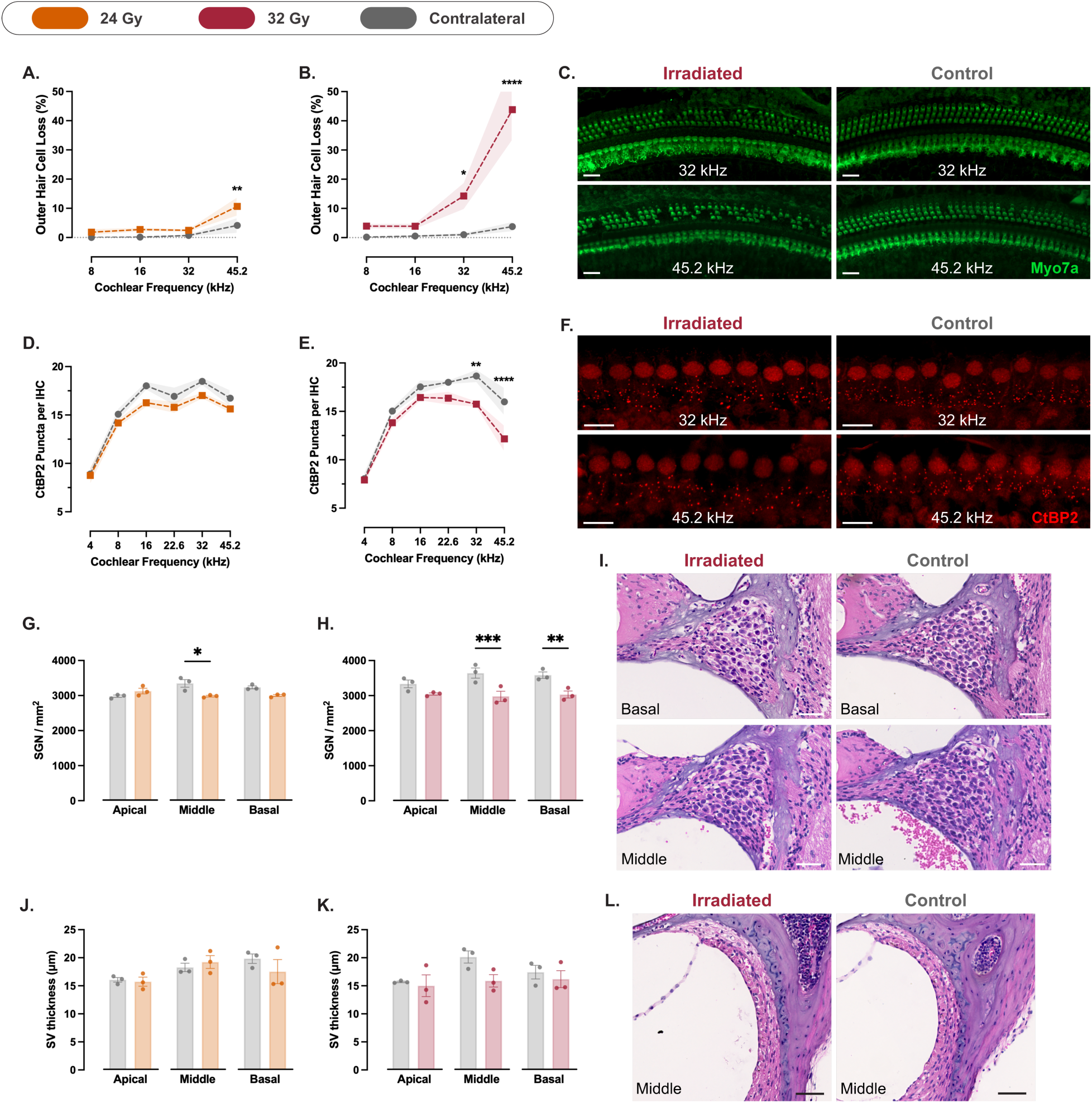
Cochlear histopathology at 4 weeks post-irradiation. (A–B) Percentage outer hair cell (OHC) loss across cochlear frequencies in mice exposed to 24 Gy (A; n = 5) and 32 Gy (B; n = 5), relative to the contralateral cochlea. OHC loss was dose-dependent and most pronounced in the 32 Gy group at high-frequency regions. (C) Representative Myo7a-stained cochlear segments from 32 Gy-irradiated and contralateral cochleae at 32 and 45.2 kHz regions. Scale bars = 20 μm. (D–E) Ribbon density measured as CtBP2-positive puncta per inner hair cell (IHC) in the 24 Gy (D; n = 5) and 32 Gy (E; n = 5) groups, showing reduced ribbon counts in the high-frequency regions of 32 Gy cochleae. (F) Representative images of CtBP2 staining at 32 and 45.2 kHz regions in irradiated and contralateral cochleae from the 32 Gy group. Scale bars = 10 μm. (G–H) Spiral ganglion neuron (SGN) density in 24 Gy (G; n = 3) and 32 Gy (H; n = 3) groups, showing reduced density at the middle turn of 24 Gy cochleae and reduced density at basal and middle turns of 32 Gy cochleae. (I) Representative hematoxylin and eosin-stained mid-modiolar cochlear sections illustrating SGN density differences between irradiated and contralateral ears at basal and middle turns from the 32 Gy group. Scale bars = 40 μm. (J–K) Stria vascularis thickness in the 24 Gy (J; n = 3) and 32 Gy (K; n = 3) groups. No significant differences were detected between irradiated and contralateral ears. (L) Representative histological sections showing the stria vascularis in irradiated and contralateral cochleae (middle turn) in the 32 Gy group. Scale bars = 50 μm. Data are presented as means with shaded areas or error bars representing SEM. Asterisks indicate statistical significance (*p < 0.05, **p < 0.01, ***p < 0.001, ****p < 0.0001). Statistical analysis was performed using two-way repeated measures ANOVA with Bonferroni correction for *post hoc* pairwise comparisons. Colors correspond to radiation dose groups as shown in the legend. The contralateral ear serves as control in all cases.

Histological analysis of the group of mice treated with the 32 Gy SRS protocol and sacrificed 16 weeks after SRS showed that IHCs were decreased by 15.6 percentage points at the 45.2 kHz region, compared to the contralateral cochlea (p = 0.04; 95% CI: 0.8 to 30.5; Fig. 5A). We also observed an OHC reduction by 48.5 percentage points at 45.2 kHz (p = 0.0003; 95% CI: 26.4 to 70.6) and 33 percentage points at 32 kHz (p = 0.005; 95% CI: 10.9 to 55.1) relative to the contralateral cochlea (Fig. 5B,C). More minor interaural differences were also observed at 16 and 8 kHz (20.1 and 15.2 percentage points, respectively), but did not reach statistical significance.

**Fig. 5.**
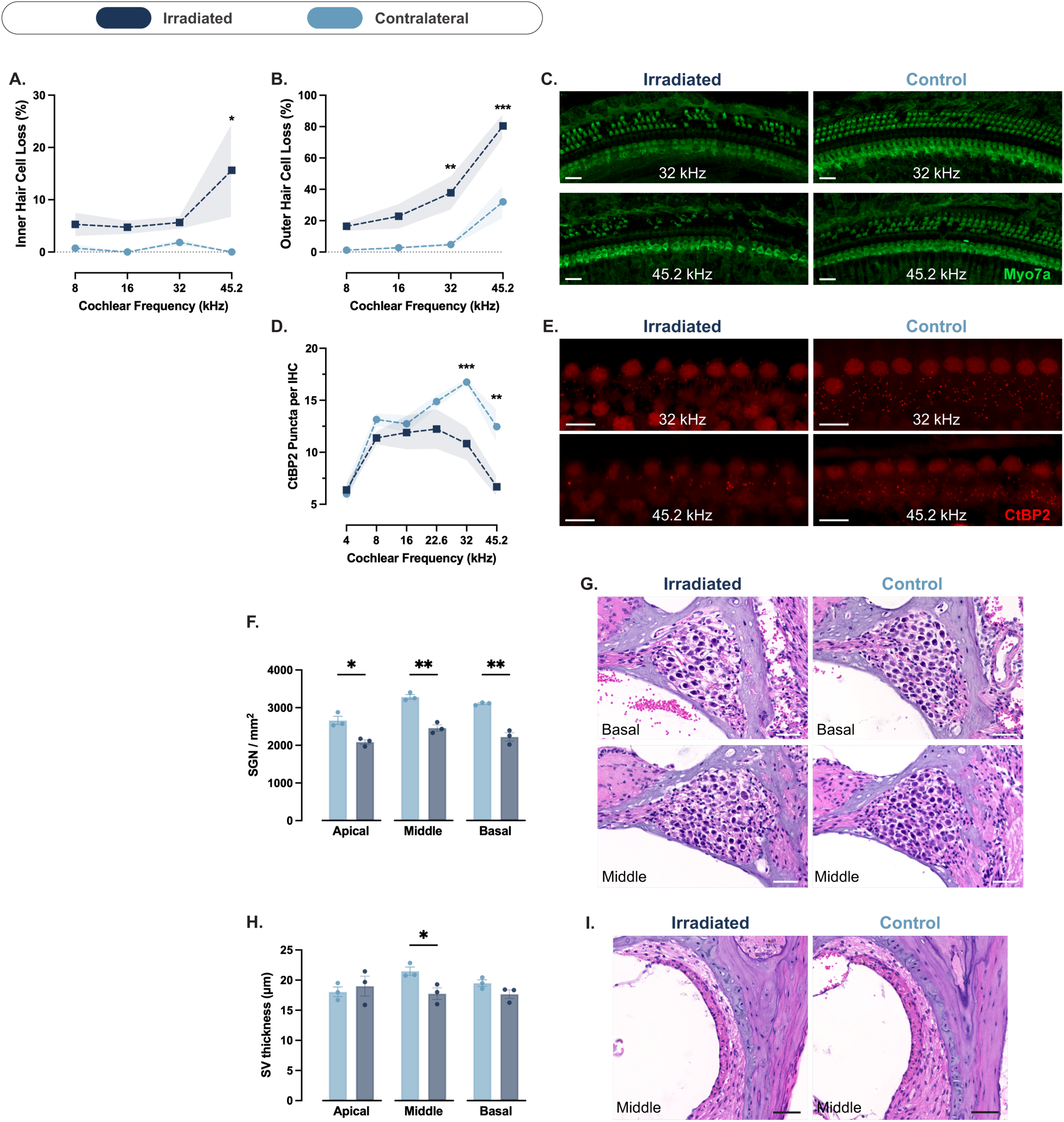
Cochlear histopathology at 16 weeks after unilateral cochlear irradiation with 32 Gy. (A–B) Percentage inner hair cell (IHC) loss (A; n = 4) and outer hair cell (OHC) loss (B; n = 4) across cochlear frequencies, relative to the contralateral cochlea. IHC and OHC loss was most pronounced at high-frequency regions. (C) Representative Myo7a-stained cochlear segments from irradiated and contralateral cochleae at the 32 and 45.2 kHz regions. Scale bars = 20 μm. (D) Ribbon density measured as CtBP2-positive puncta per IHC (n = 4), showing reduced ribbon counts in high-frequency regions of irradiated cochleae. (E) Representative images of CtBP2 staining at 32 and 45.2 kHz regions in irradiated and contralateral cochleae. Scale bars = 10 μm. (F) Spiral ganglion neuron (SGN) density (n = 3), showing reduced density at the three cochlear turns, relative to the contralateral cochlea. (G) Representative hematoxylin and eosin-stained mid-modiolar cochlear sections illustrating SGN density differences between irradiated and contralateral ears at basal and middle turns. Scale bars = 40 μm. (H) Stria vascularis thickness analysis (n = 3), demonstrating a reduced thickness at the middle turn of the irradiated cochleae compared to the contralateral side. (I) Representative histological sections showing the stria vascularis in irradiated and contralateral cochleae (middle turn). Scale bars = 50 μm. Data are presented as means with shaded areas or error bars representing SEM. Asterisks indicate statistical significance (*p < 0.05, **p < 0.01, ***p < 0.001). Statistical analysis was performed using two-way repeated measures ANOVA with Bonferroni correction for *post hoc* pairwise comparisons. Colors correspond to conditions as shown in the legend. The contralateral ear serves as control in all cases.

## 3. 5. Ribbon synapse loss is localized to high-frequency regions and exacerbated over time

We did not observe any significant reduction in the counts of ribbons per IHC in the 8, 16 (Supplementary Fig. S2K,L), and 24 Gy groups (Fig. 4D). In the 32 Gy group, compared to the contralateral cochlea, ribbons were reduced by 3.8 ribbons/IHC at 45.2 kHz (p < 0.0001; 95% CI: 1.9 to 5.7) and by 2.9 ribbons/IHC at the 32 kHz region (p = 0.001; 95% CI: 1 to 4.8; Fig. 4E,F).

At 16 weeks post-SRS, compared to the contralateral cochlea, ribbons were reduced by 5.8 ribbons/IHC at 45.2 kHz (p = 0.001; 95% CI: 2.2 to 9.3) and by 5.9 ribbons/IHC at the 32 kHz region (p = 0.0009; 95% CI: 2.4 to 9.5; Fig. 5D,E).

## 3. 6. Spiral ganglion neuron density declines in higher doses

We analyzed SGN density at week 4 in the groups treated with 24 and 32 Gy. In the 24 Gy group, SGN density was reduced in the irradiated cochlea compared to the contralateral side by 231.2 SGN/mm^2^ at the basal turn (p = 0.11; 95% CI: -66.5 to 529) and by 363.6 SGN/mm^2^ at the middle turn (p = 0.03; 95% CI: 65.8 to 661.3; Fig. 4G). In the 32 Gy group, SGN density was reduced in the irradiated cochlea compared to the contralateral side by 552.1 SGN/mm^2^ at the basal turn (p = 0.01; 95% CI: 115.7 to 988.4) and by 662.7 SGN/mm^2^ at the middle turn (p = 0.004; 95% CI: 226.3 to 1099; Fig. 4H,I).

These differences were more pronounced at 16 weeks post-SRS. Compared to the contralateral side, SGN density in the irradiated cochlea was reduced by 882.2 SGN/mm² at the basal turn (p = 0.004; 95% CI: 457.8 to 1307; Fig. 5F,G), 825.7 SGN/mm² at the middle turn (p = 0.005; 95% CI: 401.3 to 1250; Fig. 5F,G), and 574.5 SGN/mm² at the apical turn (p = 0.02; 95% CI: 150.1 to 999; Fig. 5F,G).

## 3. 7. Stria vascularis thickness is minimally affected by irradiation

We measured the thickness of the stria vascularis at week 4 in the 24 and 32 Gy groups. In the 24 Gy group, we observed no difference between the irradiated and contralateral cochleae (Fig. 4J). In the 32 Gy group, the stria vascularis was on average 21% thinner in the middle turn of the irradiated cochlea compared to the contralateral side (mean difference, 4.3 μm; Fig. 4K,L). However, this difference failed to reach statistical significance (p = 0.2). There was no difference between the two sides at the basal and apical turns.

At 16 weeks post-SRS, the interaural difference in stria vascularis thickness at the middle turn did not increase compared to week 4. However, due to reduced variance in the measurements, the difference reached statistical significance (mean difference, 3.7 µm; p = 0.02; 95% CI: 0.7 to 6.7; Fig. 5H,I). No significant differences were observed at the basal or apical turns.

## 4. Discussion

### 4.1. Summary of key findings

In this study, we established a mouse model of SRS-induced unilateral hearing loss using a near-cochlear target with a clinical LGK device. The irradiation protocol was well tolerated, with no systemic side effects observed, highlighting the 3R compatibility of the new model. We observed a reproducible, dose-dependent, predominantly high-frequency, and progressive hearing loss on the irradiated side. Histological analysis corroborated a corresponding decrease in OHC survival, ribbon density, and SGN density, particularly in the basal and middle cochlear turns. The 32 Gy protocol induced consistent and measurable cochlear toxicity. The contralateral cochlea remained unaffected, validating its use as an internal control.

### 4.2. Comparison with previous models

In previous animal studies, researchers employed a variety of irradiators such as Cobalt-60 teletherapy units (10, 12), cabinet X-ray irradiators (8, 9, 13, 18), and clinical linear accelerators (7, 11, 14–16), which delivered broad-field or regional irradiation without focal stereotactic guidance. These devices have been used to induce hearing loss, investigate the radiobiological effects of radiation, and evaluate potential otoprotective strategies. For this purpose, authors used a wide range of radiation doses in a single fraction to the cochlea, ranging from 10 to 60 Gy, with varying functional and histological outcomes. These setups lacked the spatial precision of stereotactic systems and often exposed both cochleae and central auditory structures to radiation. Our experimental setting, employing an image-guided approach and clinical SRS equipment, enables true 3D stereotactic targeting of small, deep structures, such as the cochlea, with submillimeter precision, steep dose gradients, and minimal off-target exposure, mirroring human SRS in VS treatment. The ability to selectively irradiate one cochlea while largely sparing the contralateral side provides an internal control, thereby reducing the number of experimental animals and inter-individual variability. This is also more clinically relevant, as SRS may be less toxic to the central auditory pathways, thereby minimizing its impact on central ABR wave components and enabling more specific investigation of peripheral neurosensory effects.

### 4.3. Radiation dose optimization and relevance to clinical settings

To develop a mouse model that allows for the quantitative assessment of radiotoxic effects, we tested four doses of irradiation: 8, 16, 24, and 32 Gy, delivered in a single fraction. The two lower doses induced only a mild OHC loss at the 45.2 kHz region and no ABR shifts in frequencies up to 32 kHz at week 4 post-SRS. The 24 Gy protocol resulted in mild ABR shifts, with a greater effect at high frequencies, as well as mild OHC and SGN loss. These effects were more pronounced and consistent in the 32 Gy protocol, resulting in both high-frequency ABR shifts and histological damage (OHC, ribbon loss, and SGN reduction); therefore, we conclude that 32 Gy is required for robust and measurable cochlear toxicity at week 4. This dose exceeds the 12 Gy marginal dose typically prescribed in human SRS for VS, which results in a cochlear dose of 12 Gy or often lower. Several factors may account for this discrepancy. First, interspecies differences in cochlear radiosensitivity could contribute, with murine cochlear tissue potentially exhibiting greater radioresistance than that of humans. Indeed, smaller animals, such as mice, are generally more radioresistant, as evidenced by higher LD_50_ values (the radiation dose causing 50% mortality) following total body irradiation (24). Second, the markedly smaller size of the mouse cochlea, approximately one-hundredth the volume of the human cochlea (25, 26), alters dose distribution and may necessitate higher absolute doses to achieve equivalent biological effects (27). Third, our model was designed to produce detectable hearing loss within a relatively short timeframe (4 weeks post-SRS) to minimize confounding by age-related hearing loss, whereas in humans, radiation-induced auditory decline often develops gradually over months or years (6). Indeed, in our long-term follow-up group, we observed a progressive decrease in hearing, as well as greater interaural histological differences, at week 16 post-SRS. Finally, our model does not include a VS. In patients, cochlear damage may be exacerbated by tumor-derived proinflammatory cytokines, such as TNF-α, which compromise cochlear integrity even prior to SRS and may synergize with the radiation-induced inflammatory response (28, 29).

### 4.4. Basal turn vulnerability

We observed a higher vulnerability of the basal turn of the cochlea to irradiation. In line with this, previous animal studies have found a higher basal turn vulnerability to irradiation (7, 8, 11, 18). Interestingly, the same hearing loss pattern is observed in patients undergoing irradiation (30). Hoistad et al., in their histopathological study of human temporal bones, corroborated this observation, finding more marked histological damage (SGN, IHC, and OHC) in the basal turn of patients treated with irradiation for head and neck tumors (31). These results are in accordance with the heightened susceptibility of the basal turn to acquired damage across multiple etiologies, including ototoxic drugs, noise exposure, and aging (32–34). This vulnerability can stem from the cochlea’s inherent capacity to generate and handle oxidative stress. Genes related to oxidative stress were found to be upregulated in the basal turn (35). Moreover, our group has previously demonstrated that NADPH oxidase 3 (Nox3), an enzyme producing reactive oxygen species (ROS), follows a tonotopic pattern of induced expression after noise trauma (33). Adding to that, Sha et al. found that basal turn OHCs had a lower level of the inherent antioxidant glutathione (36). In the context of irradiation, it is well established that most of the cellular damage results from the indirect effects of irradiation and is mediated by ROS (37). Based on the above, we can hypothesize that basal vulnerability may be due to excessive and sustained radiation-induced ROS production, combined with a limited antioxidant capacity.

### 4.5. Progressive and irreversible hearing loss

Hearing loss was irreversible and progressive up to 16 weeks post-SRS. In our 32 Gy experimental group, followed for 16 weeks, we observed a progressive increase in threshold shifts, which largely outweighed the natural age-related changes seen in the contralateral ear. This was also corroborated by the progressive decline in wave I amplitudes. Moreover, the histological damage induced by irradiation at 16 weeks exceeded that observed at 4 weeks. This progressive decline was observed in previous animal studies with large-field irradiation protocols (9, 12, 13). Liu et al. found a progressive increase in ABR thresholds and a progressive decline in wave I amplitudes from 7 to 21 days after 20 Gy irradiation of C57BL/6J mice (12). Ribbon density and size were also reduced progressively during this period. In another study, Gao and colleagues describe a progressive OHC loss and stria vascularis atrophy from 4 to 8 weeks using a 15 Gy protocol (18). Mujica-Mota and colleagues treated guinea pigs with a fractionated protocol (total dose 70 Gy, 3.5 Gy per fraction) and observed progressively increasing ABR thresholds at 1, 6, and 16 weeks (13). This progressive hearing loss pattern is also seen in clinical studies. Indeed, an irreversible and progressive hearing loss is observed in patients who undergo SRS for a VS or radiation therapy for head and neck tumors (6, 30). This pattern has also been observed after different initial insults. Both clinical and animal studies have shown an acceleration of hearing loss over time following noise trauma (38, 39). The underlying mechanisms comprise oxidative damage and vascular dysfunction, leading to the loss of SGN and ribbon synapses (40).

### 4.6. Limitations and future directions

While this animal model introduces several improvements over earlier versions, there are still multiple areas for further refinement and expansion. The irradiation protocol proposed in this study was tested in one mouse strain (C57BL/6J), and thus, extrapolating the exact functional and histological outcomes for other mouse strains or animal models must be done with caution. C57BL/6J mice are known to develop progressive hearing loss as early as 12 weeks of age due to the age-related hearing loss allele of the cadherin 23 gene (41). In our study, we irradiated 6-week-old mice and observed progressive threshold shifts at the contralateral ear, specifically at 32 kHz, consistent with previous studies (42). Using the contralateral ear as an internal control, we were able to overcome this limitation and isolate the effect of radiation treatment. Moreover, in our model, mice do not bear a VS and thus differ from the exact clinical setting. Injection of schwannoma cells into the cerebellopontine angle of the mouse brain can be performed, leading to the formation of a VS (43). In the current study, we aimed to isolate the effect of radiation independent of the presence of a VS. Future studies can combine the development of a VS with targeted SRS. In this case, special consideration should be given to the size and distance of the VS to the cochlea, which can greatly influence the dose delivered to the cochlea. We performed histological analysis at two time points (4 and 16 weeks) but did not include intermediate time points that could reveal the temporal evolution of radiation-induced cochlear pathology. Such data could improve understanding of damage progression. Another limitation of our study is the lack of evaluation of the vestibular system. We focused on characterizing hearing loss, but SRS can also exacerbate vestibular symptoms in treated patients (5, 44). Although we did not observe an apparent vestibular phenotype in our mouse model, detailed vestibular testing merits further investigation.

### 4.7. Conclusion

In conclusion, we present a clinically relevant and technically robust mouse model of radiation-induced unilateral hearing loss using an SRS device. This model reliably reproduces key features of cochlear toxicity observed in patients undergoing SRS for VS, including dose-dependent, progressive, and high-frequency hearing loss accompanied by OHC, ribbon synapse, and SGN loss. Importantly, the use of image-guided targeting not only ensures reproducibility and anatomical specificity but also avoids the systemic side effects reported in previous broad-field irradiation models. The inclusion of an internal contralateral control further reduces variability and animal use. Together, these features make this model a valuable platform for investigating the radiobiological mechanisms of radiation-induced cochlear injury and for evaluating candidate radioprotective strategies.

## Author contributions

**Dimitrios Daskalou**: Conceptualization, Methodology, Data curation, Investigation, Formal analysis, Project administration, Validation, Visualization, Writing – original draft, Writing – review & editing. **Francis Rousset**: Conceptualization, Methodology, Investigation, Formal analysis, Resources, Visualization, Validation, Supervision, Writing – original draft, Writing – review & editing. **Stéphanie Sgroi**: Investigation, Validation, Writing – review & editing. **Lucie Oberhauser**: Investigation, Writing – review & editing. **Jean-Philippe Thiran**: Conceptualization, Methodology, Writing – review & editing. **Constantin Tuleasca**: Methodology, Writing – review & editing. **Ileana O Jelescu**: Conceptualization, Methodology, Writing – review & editing. **Marc Levivier**: Conceptualization, Methodology, Investigation, Formal analysis, Validation, Resources, Writing – review & editing, Supervision. **Pascal Senn**: Conceptualization, Methodology, Formal analysis, Validation, Resources, Writing – review & editing, Supervision.

## Funding

The Swiss Academy of Medical Sciences MD–PhD grant (323630_214546) supported D.D. IOJ is supported by an Eccellenza fellowship from the Swiss National Science Foundation (194260). A Swiss National Science Foundation grant supports FR and PS (204546).

## Acknowledgments

We gratefully acknowledge Nicolas Liaudet and Sergei Startchik from the Bioimaging Core Facility of the University of Geneva for their expert support in imaging analysis. We also thank Cristian Cotrutz from the Lausanne Gamma Knife Center for his valuable assistance in developing the stereotactic irradiation protocol, and Nicolas Dupuy for his help during animal experimentation at the same facility.

## Conflicts of interest

The authors disclose no conflicts of interest.

## Data availability

Data will be made available on request.

## Notes

### Competing Interest Statement

The authors have declared no competing interest.

