## Supplementary figures for "Introducing a translationally relevant mouse model of radiosurgery-induced unilateral hearing loss"

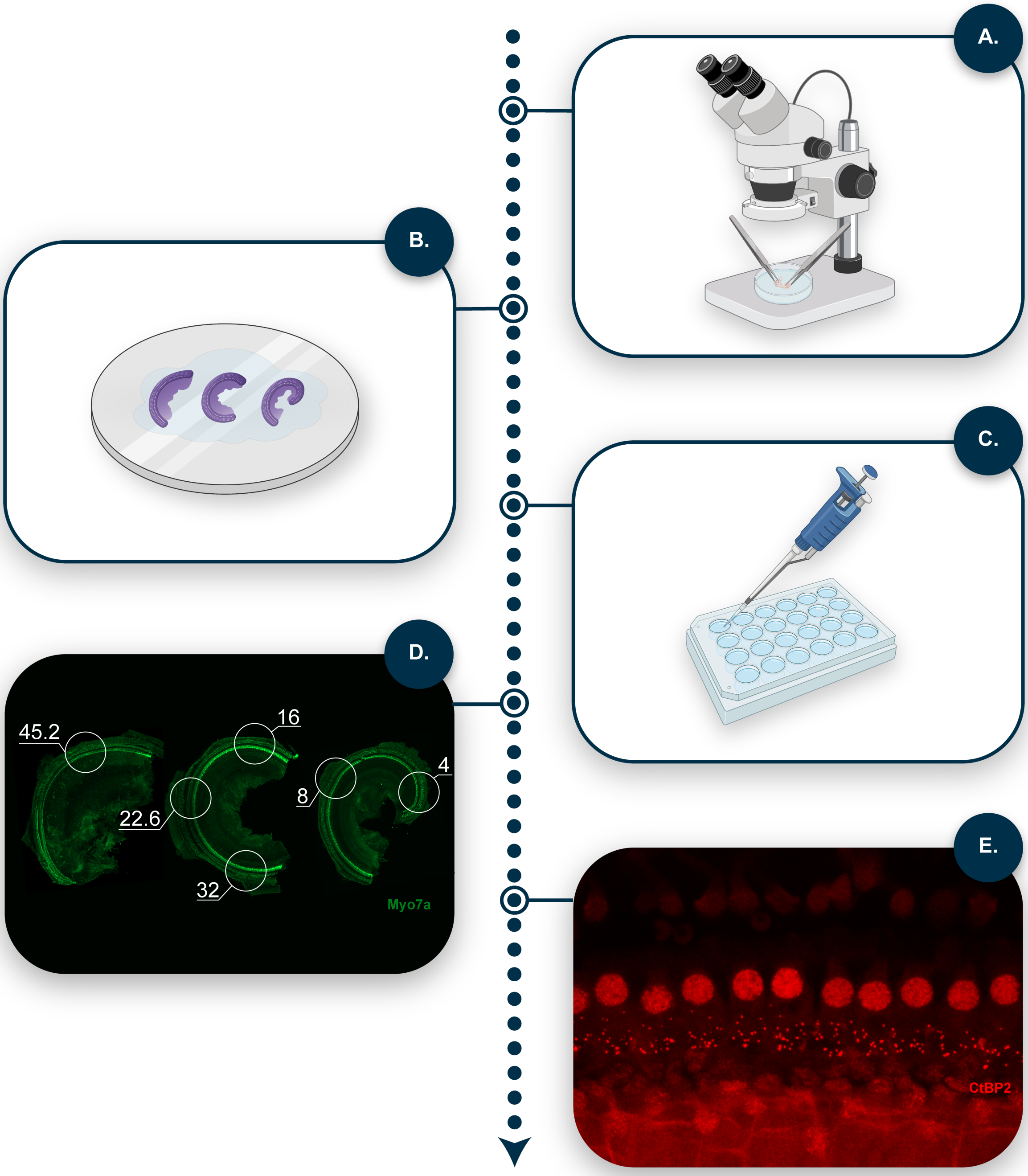


**Supplementary Fig. S1 Cochlear dissection, whole-mount preparation, and frequency-specific quantification workflow.**
(A) Dissection of the decalcified cochlea and segmentation of the cochlear sensory epithelium into apical, middle, and basal turns under a stereomicroscope.
(B) Adhesion of cochlear segments onto 12 mm round coverslips using Cell-Tak tissue adhesive.

(C) Coverslips with samples were placed in PBS in 24-well plates and processed for immunostaining.
(D) Representative cytocochleogram showing cochlear frequency mapping generated using the Measure_Line plugin from the Eaton-Peabody Laboratories Histology Core (Mass Eye and Ear, https://masseyeandear.org/research/otolaryngology/eaton-peabody-laboratories/histology-core), enabling quantification of hair cell survival in 300 μm-long segments at defined frequency regions.
(E) The previously generated tonotopic map was used as a guide to acquire high-magnification confocal images of the inner hair cell region, within 10–15 inner hair cells at each frequency region.
Panels A–C include elements created with BioRender.com.

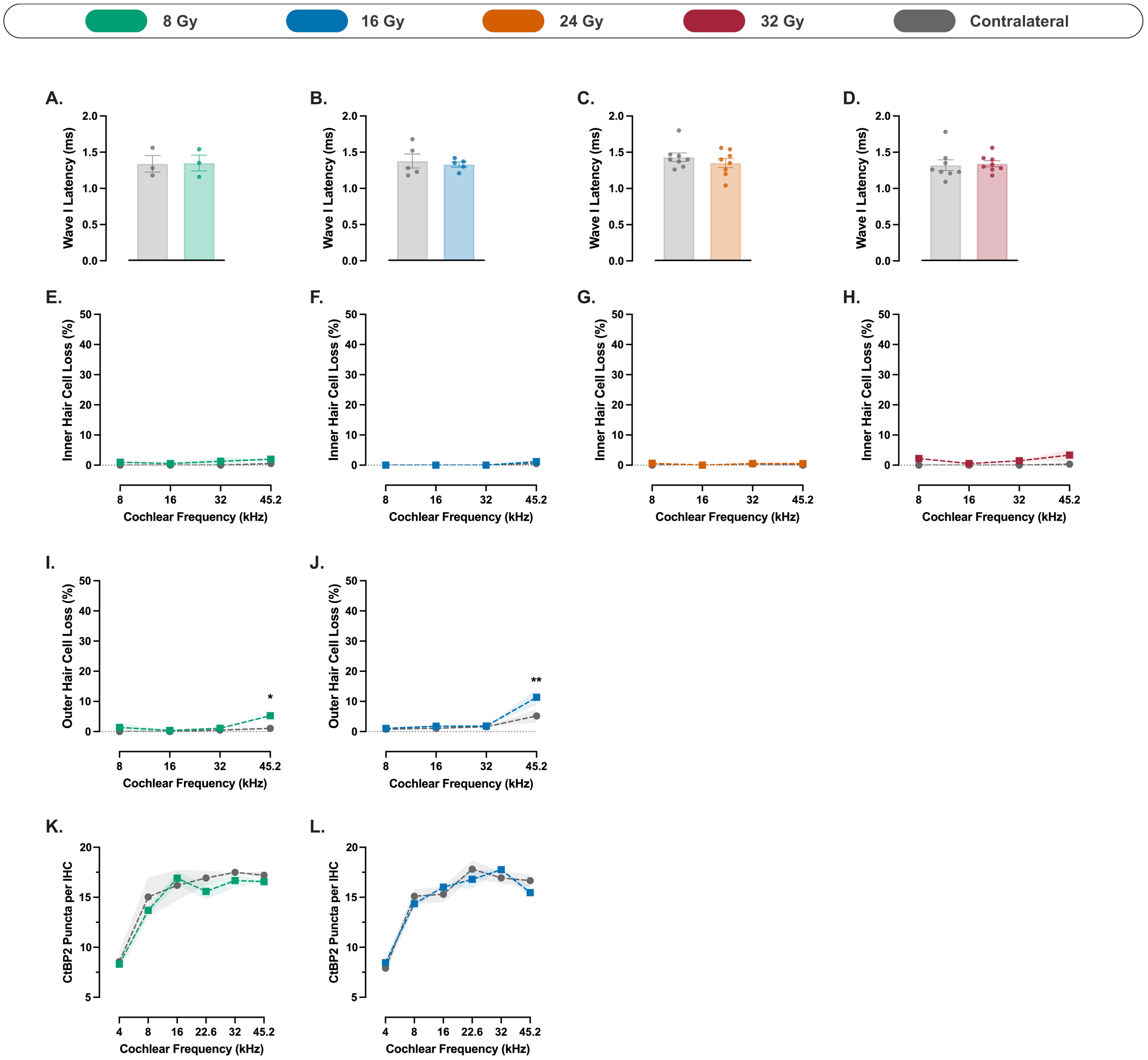


**Supplementary Fig. S2 Additional auditory and histological outcomes across radiation dose groups.**

(A–D) Wave I latency at week 4 following cochlear irradiation with 8 Gy (A; n = 3), 16 Gy (B; n = 5), 24 Gy (C; n = 8), and 32 Gy (D; n = 8), showing no significant interaural differences.

(E–H) Inner hair cell (IHC) loss across cochlear frequencies at week 4 in the 8 Gy (E; n = 3), 16 Gy (F; n = 5), 24 Gy (G; n = 5), and 32 Gy (H; n = 5) groups, with no significant differences observed between irradiated and contralateral ears.

(I–J) Percentage outer hair cell (OHC) loss across cochlear frequencies in mice exposed to 8 Gy (I; n = 3) and 16 Gy (J; n = 5), relative to the contralateral cochlea, showing mild but significant interaural differences at 45.2 kHz.

(K–L) Ribbon synapse density measured as CtBP2-positive puncta per inner hair cell in the 8 Gy (K; n = 3) and 16 Gy (L; n = 5) groups, showing no significant differences between irradiated and contralateral cochlea.

Data are presented as means with error bars or shaded areas representing SEM. Asterisks indicate statistical significance (*$p < 0.05$, **$p < 0.01$). Statistical analysis was performed using two-way repeated measures ANOVA with Bonferroni correction for *post hoc* pairwise comparisons. Colors correspond to radiation dose groups as shown in the legend. The contralateral ear serves as control in all cases.
